# New evidence concerning the genome designations of the AC(DC) tetraploid *Avena* species

**DOI:** 10.1101/2020.10.02.323345

**Authors:** Honghai Yan, Zichao Ren, Di Deng, Kehan Yang, Chuang Yang, Pingping Zhou, Charlene Wight, Changzhong Ren, Yuanying Peng

**Affiliations:** Triticeae Research Institute, Sichuan Agricultural University, Chengdu, 611130, China; Agriculture and Agri-Food Canada, Ottawa Research and Development Centre, 960 Carling Ave, Ottawa ON K1A0C6 Canada; Baicheng Academy of Agricultural Sciences, Baicheng, 137000, China

## Abstract

The tetraploid *Avena* species in the section *Pachycarpa* Baum, including *A. insularis, A. maroccana*, and *A. murphyi*, are thought to be involved in the evolution of hexaploid oats; however, their genome designations are still being debated. Repetitive DNA sequences play an important role in genome structuring and evolution, so understanding the chromosomal organization and distribution of these sequences in *Avena* species could provide valuable information concerning genome evolution in this genus. In this study, the chromosomal organizations and distributions of six repetitive DNA sequences (including three SSR motifs (TTC, AAC, CAG), one 5S rRNA gene fragment, and two oat A and C genome specific repeats) were investigated using non-denaturing fluorescence in situ hybridization (ND-FISH) in the three tetraploid species mentioned above and in two hexaploid oat species. Preferential distribution of the SSRs in centromeric regions was seen in the A and D genomes, whereas few signals were detected in the C genomes. Some intergenomic translocations were observed in the tetraploids; such translocations were also detected between the C and D genomes in the hexaploids. These results provide robust evidence for the presence of the D genome in all three tetraploids, strongly suggesting that the genomic constitution of these species is DC and not AC, as had been thought previously.

## Introduction

The cultivated hexaploid oat, *Avena sativa* L. (2n=6x=42, genomes AACCDD), is the sixth most important cereal crop cultivated worldwide [1]. The superior nutraceutical properties of the oat grain have attracted considerable attention from both breeders and consumers [2]. The genus *Avena* L. comprises a number of closely related species with a basic chromosome number of seven, and includes diploids, tetraploids, and hexaploids [3]. *A. sativa* is an allohexaploid species displaying disomic inheritance, and is closely related to the weedy species *A. sterilis* L. [4]. It is believed that *A. sativa* was domesticated from *A. sterilis* somewhere in Northwest Asia [4, 5].

The evolutionary history of the hexaploid oat A, C, and D genomes has been under intense scrutiny [6–13]. The identities of its genome donors, however, remain inconclusive. It has been assumed that one of the species in the section *Pachycarpa* Baum, which includes *A. insularis* Ladiz., *A. maroccana* Gdgr. (synonym, *A. magna* Murphyi et Terr.), and *A. murphyi* Ladiz., has been involved in the formation of the hexaploids [13–15]. Currently, these tetraploid species have all been designated as having AC genomes (2n=4x=28, genomes AACC). The most challenging mystery has concerned the origin of the D genome donor, since no natural diploid with a D genome has been identified. Because of the high homology between the A and D genomes in *A. sativa* [9], the D genome in the hexaploid has been considered to be a variant of the A genome, suggesting that it originated from one of the A genome diploids [9, 16]. Furthermore, the AC genome designations of the three tetraploids in the section *Pachyacarpa* have been challenged by evidence from both cytogenetic [17] and molecular studies [13, 18]. Fluorescence in situ hybridization (FISH) with an A genome-specific probe did not produce hybridization signals in the chromosomes of the AC genome tetraploids, and this absence was also observed in the D and C genome chromosomes of the hexaploids [17]. Another previous study, which used high-density genotyping-by-sequencing (GBS) markers, also showed that the hexaploid D genome, rather than the hexaploid A genome, has stronger matches with the A genome of these tetraploids [13]. Thus, more evidence is needed to confirm the genomic composition of these tetraploids.

Repetitive DNA elements are major components of plant genomes, including those of the *Avena* species, where more than 70% of the genome is predominated by repetitive DNA sequences [19]. Generally, repetitive DNA sequences evolve more rapidly than genic sequences, and play essential roles in genome structuring and evolution [20]. Their organization, distribution, and density can be specific for a species, a genome, or even a chromosome [21]. Therefore, extensive examination of the organization and distribution of repetitive DNA sequences could assist our understanding of the organization and evolution of genomes [22, 23], provide valuable information in taxonomic and phylogenetic studies [24], and help develop diagnostic markers for identifying specific chromosomes and chromosome regions [25–27].

Fluorescence in situ hybridization (FISH) is one of the most routinely-used tools to study the physical organization and distribution of repetitive DNA sequences. Indeed, FISH techniques using repetitive DNA sequences have been shown to be powerful tools in cytogenetic studies of *Avena* species. For instance, FISH using oat A and C genome-specific repetitive DNA sequences as probes clearly differentiated the A, C, and D genomes of hexaploid oat [17]. However, conventional FISH analysis is time-consuming because of the preparation and labeling of probe sequences and the denaturing of probes and chromosomes [27]. In recent years, a new FISH labeling procedure, non-denaturing FISH (ND-FISH), has been developed. ND-FISH uses synthesized oligonucleotide sequences as probes, and performs FISH analysis under non-denaturing conditions, thus substantially simplifying the procedure [28]. It has been widely used for cytogenetic studies in barley [25, 29], wheat [26], and rye [26], but less often for oat [30–33].

In this study, we used ND-FISH analysis to analyze the chromosomal organization of three tri-nucleotide SSR motifs (TTC, AAC, and CAG) and three oligonucleotide sequences (oligo-Am1, oligo-As120a, and oligo-Ta794) in *Avena* spp. The latter probes were derived from the oat A and C genome-specific repetitive DNA sequences Am1 and As120a and the wheat 5S rRNA gene. The relationships amongst the genomes from three tetraploid species in the section *Pachycarpa*, as well as two hexaploid oat species, were determined.

## Materials and Methods

### Plant materials

Table 1 shows the plant materials used in this study, which comprised two hexaploid species (*A. sativa* and *A. sterilis*) and three AC(DC) genome tetraploid species (*A. insularis*, *A. marrocana*, and *A. murphyi*). Seeds of these materials were obtained from either the United States Department of Agriculture (USDA), or Plant Gene Resources of Canada (PGRC), with the exception of *A. insularis*, for which the seeds were kindly provided by Dr. Rick Jellen, Brigham Young University, Provo, UT, USA.

**Table 1.**
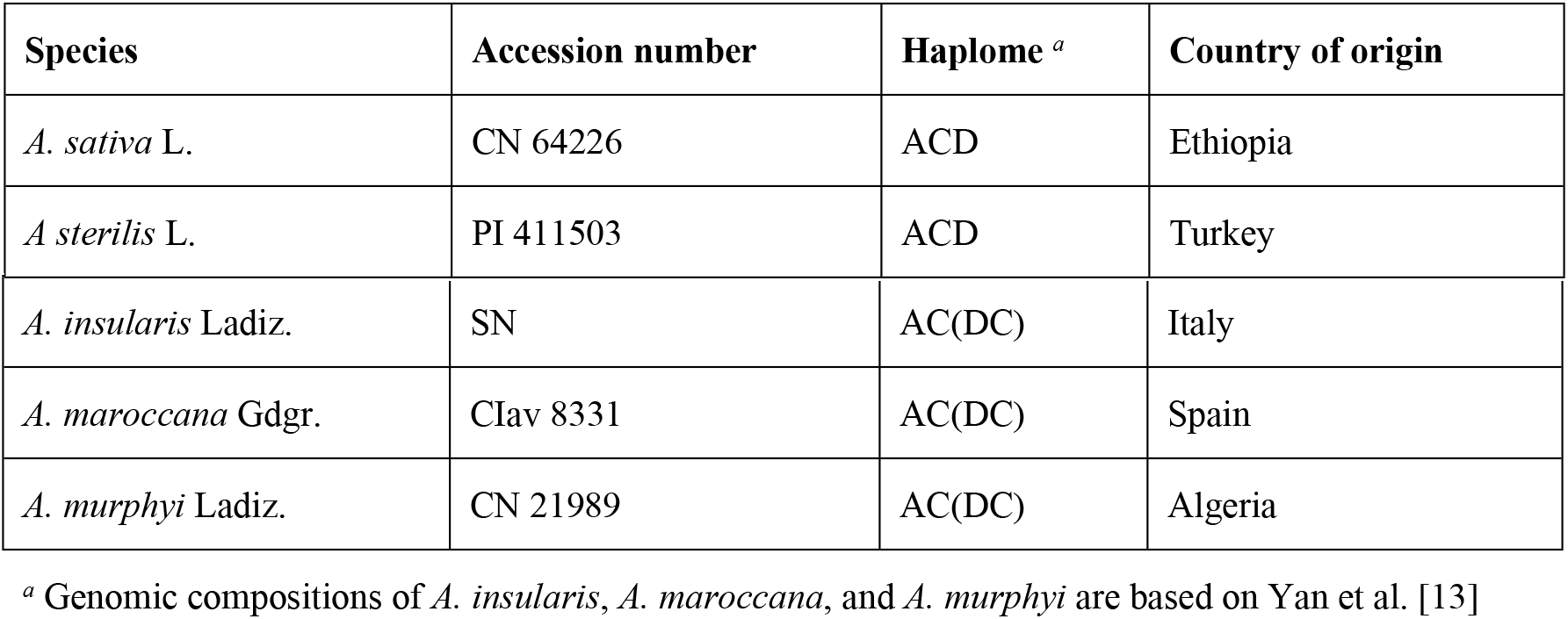
List of materials used in this study, including species name, accession number, haplome and origin.

| Species | Accession number | Haplome <sup>a</sup> | Country of origin |
| --- | --- | --- | --- |
| *A. sativa* L. | CN 64226 | ACD | Ethiopia |
| *A. sterilis* L. | PI 411503 | ACD | Turkey |
| *A. insularis* Ladiz. | SN | AC(DC) | Italy |
| *A. maroccana* Gdgr. | Clav 8331 | AC(DC) | Spain |
| *A. murphyi* Ladiz. | CN 21989 | AC(DC) | Algeria |

<sup>a</sup> Genomic compositions of *A. insularis*, *A. maroccana*, and *A. murphyi* are based on Yan et al. [13]

### ND-FISH probes

Three SSR motifs ((TTC)_5_, (AAC)_5_, (CAG)_5_), two oligonucleotides derived from oat A and C genome specific repeats, and a wheat 5S rRNA gene fragment were used as ND-FISH probes. TTC, AAC, and CAG are tri-nucleotides that are highly abundant and widely distributed throughout the barley, wheat, and rye genomes [34, 35]. The other three probes are: (1) oligo-As120a, a 71bp fragment specific to the oat A genome that was isolated from *A. strigosa* [17]; (2) oligo-Am1, a 51bp fragment specific to the oat C genome that was isolated from *A. murphyi* [36]; and (3) oligo-Ta794, a 41bp 5S rRNA sequence fragment that was isolated from *T. aestivum* [37]. All of these probes were synthesized by Sangon Biotech Co., Ltd. (Shanghai, China). The synthesized oligonucleotides were 5’-end labeled with either 6-carboxytetramethylrhodamine (TAMRA) or 6-carboxyfluorescein (FAM). Further details concerning the probes, including their names, DNA sequences, and fluorochromes, are listed in Table 2.

**Table 2.**
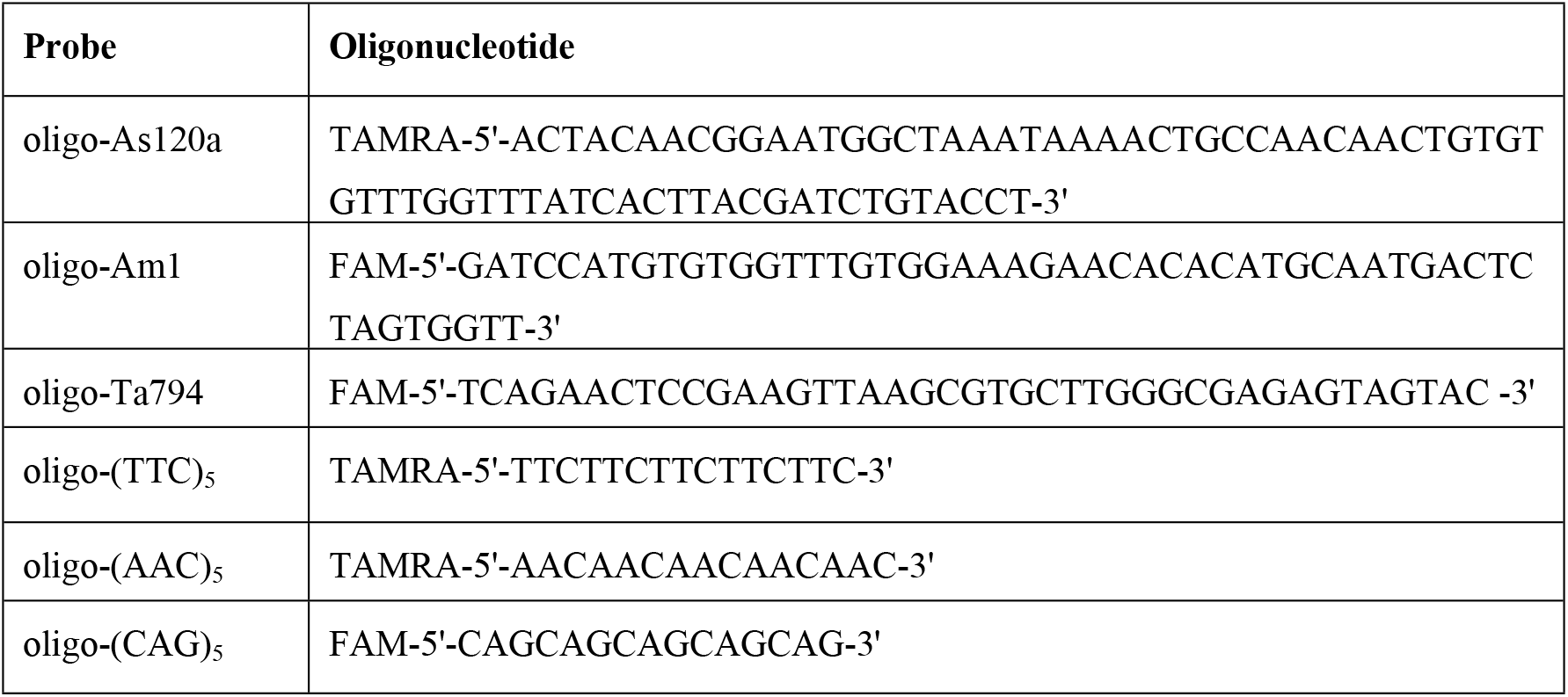
Oligonucleotide probes used for fluorescence in situ hybridization (FISH) analysis

### Preparation of metaphase spreads

Metaphase chromosome preparation paralleled that of previous experiments with some modifications [33]. In brief, seeds were imbibed in distilled water for 18 h at 25°C in the dark, and then placed in petri dishes lined with a layer of moist filter paper. To synchronize cell division and accumulate metaphase plates, the germinated seeds were transferred to a 4°C growth cabinet for 48 h, then to one at 25°C for 24 h. Root tips were harvested from germinated seeds, pre-treated in 1.0 MPa nitrous oxide gas for 3.5 h, then fixed in glacial acetic acid for 5 to 20 min before being fixed in 70% ethanol. Apical meristems were extruded from the fixed root tips and enzymatically digested with 2% cellulose and 1% pectinase. After being squashed in a drop of 60% acetic acid, each suspension was dropped onto a clean glass slide. The slides were air-dried prior to ND-FISH analysis.

### ND-FISH analysis

ND-FISH was performed as described by Fu et al. [26]. Briefly, air-dried, pre-treated slides were fixed for 10 min with 4% (w/v) paraformaldehyde, and then immersed in 2×saline sodium citrate (SSC) for 10 min. After dehydration in an ice-cold ethanol series of 75%, 95%, and 100% for 5 min each, they were air-dried These air-dried slides were then denatured at 80°C for 2 min in deionized formamide (60 μL per slide), followed by dehydration in 75%, 95%, and 100% alcohol at −20°C for 5 min each before air drying. A 10 μL aliquot of a hybridization mixture containing 0.5 μL FISH probe, 4.75 μL 2×SSC, and 4.75 μL 1×TE was applied to each slide, then the slides were incubated for 2 h at 37°C. The slides were counterstained with DAPI and Vectashield mounting medium (Vector Laboratories, Inc., Burlingame, CA, USA). Sequential FISH analyses were performed as described in Fominaya et al. [33]. Digital images were captured using an Olympus BX-51 epifluorescence microscope equipped with a Photometric SenSys Olympus DP70 CCD camera (Olympus, Tokyo), and processed using Photoshop V7.0 (Adobe Systems Incorporated, San Jose, CA). After capturing each image, the slides were washed as described by Komuro et al. [35].

## Results

The assignments of each chromosome of the tetraploid and hexaploid oat lines were based on the work of Fominaya et al. [33]. The A and C genome-specific probes oligo-Am1 and olligo-As120a, as well as the 5S rRNA probe oligo-pTa794, were used to assist with chromosome identification, enabling the A and C genome chromosomes to be distinguished from the D genome chromosomes.

### Chromosomal organization of repeats in three tetraploid species

As expected, the oligo-Am1 probe produced multiple strong signals all along half of the chromosomes in all three tetraploids (Figs 1b, 1h and 1n). These chromosomes were identified as being the C genome chromosomes, meaning the remaining chromosomes should belong to the A(D) genome. Six C/A(D) translocations were observed in *A. insularis* (Fig 1b and 1s) and *A. maroccana* (Figs 1h and 1s), while only four were found in *A. murphyi* (Figs 1n and 1s). The oligo-Ta794 probe produced two pairs of bright signals on chromosome 1 A(D) in *A. insularis* (Figs 1c and 1s) and *A. maroccana* (Figs 1i and 1s), and on chromosome 4A(D) in *A. murphyi* (Figs 1o and 1s). In addition, a pair of weak signals was detected on chromosome 2C of *A. insularis* (Figs 1c and 1s). No discernable hybridization signals were detected in any of the three tetraploids using the oligo-As120a probe (Figs 1a, 1g and 1m).

**Fig 1.**
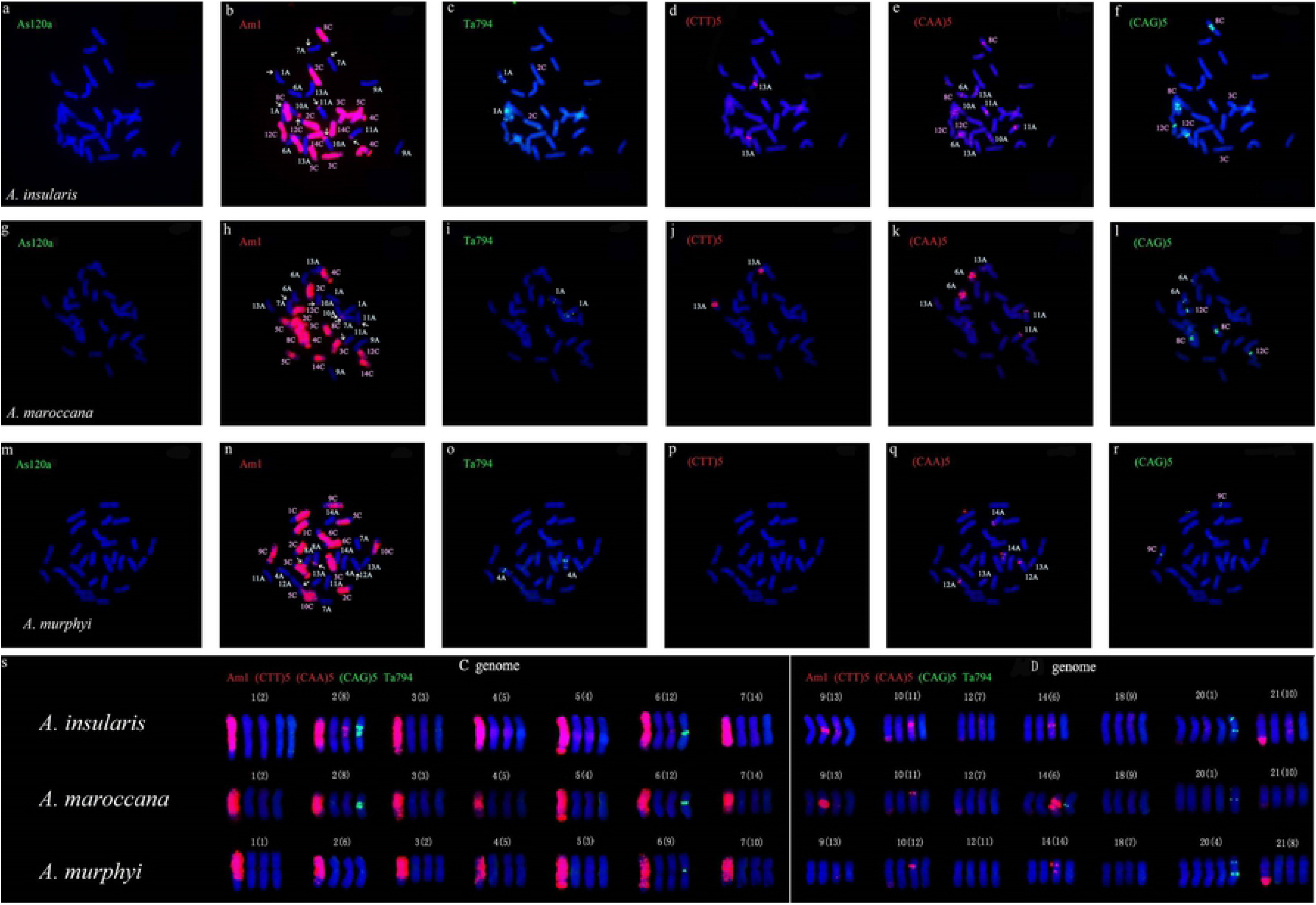
Karyotypes of the tetraploid species *A. insularis* (a-f), *A. maroccana* (g-l), and *A. murphyi* (m-r) after sequential FISH analysis. The probes included FAM-As120a (green), TAMRA-Am1 (red), FAM-Ta794 (green), TAMRA-(TTC)_5_ (red), TAMRA-(AAC)_5_ (red), and FAM-(CAG)_5_ (green). (s) Karyotype of single metaphase chromosomes of these species. The white arrows indicate the intergenomic translocations.

For the SSR probes, the oligo-(TTC)_5_ probe produced strong signals on chromosome 13A(D) in *A. insularis* (Figs 1d and 1s) and *A. maroccana* (Figs 1j and 1s), covering a large region around the centromeres. However, no such signals were detected in *A. murphyi* (Figs 1p and 1s). Many more signals were produced by oligo-(AAC)_5_ than by oligo-(TTC)_5_. In *A. insularis*, oligo-(AAC)_5_ produced strong signals on chromosomes 8C, 6A(D), 10 A(D), 11 A(D), and13 A(D), with faint signals on 12C and the remaining three A(D) genome chromosome pairs. These signals were found in centromeric regions, intercalary regions, or both (Figs 1e and 1s). In the other two tetraploids, chromosome pairs with (AAC)_5_ hybridization signals were reduced to three, all belonging to the A(D) genome, but the signal patterns were highly differentiated both in distribution and intensity. In *A. maroccana*, signals on 6A(D) were very strong and covered a large region around the centromere, while signals on 11A(D) and 13A(D) were weak and located in the centromeric and telomeric regions, respectively (Figs 1k and 1s). In *A. murphyi*, signals on 12A(D) were positioned in the centromeric region, while signals on 14A(D) were located in the centromeric region and on the short arm. Very weak signals on the long arm were observed on 13A(D) (Figs 1q and 1s). The oligo-(CAG)_5_ probe produced signals on two (8C and 12C) (Figs 1f and 1s), three (8C, 12C and 6A(D)) (Figs 1l and 1s) and one (9C) chromosome (Figs 1r and 1s) in *A. insularis, A. maroccana*, and *A. murphyi*, respectively. All of these signals were located in centromeric regions, but differed in intensity.

### Chromosomal organization of repeats in two hexaploid species

In the two hexaploids, the oligo-Am1 and oligo-As120a probes produced multiple signals that were evenly distributed along 14 chromosomes each, identifying these chromosomes as belonging to the the C and A genomes, while indicating that the remaining 14 chromosomes belong to the D genome (Figs 2a and 2f). Three minor C/D translocations, on chromosomes 10D, 20D, and 21D, were detected in *A. sativa* (Figs 2a and 2j), whereas two minor C/D translocations, on 10D and 21D, were observed in *A. sterilis* (Fig 2f and 2j). Two 5S sites were detected in A. sativa, on chromosomes 19A and 20D (Figs 2b and 2j).

**Fig 2.**
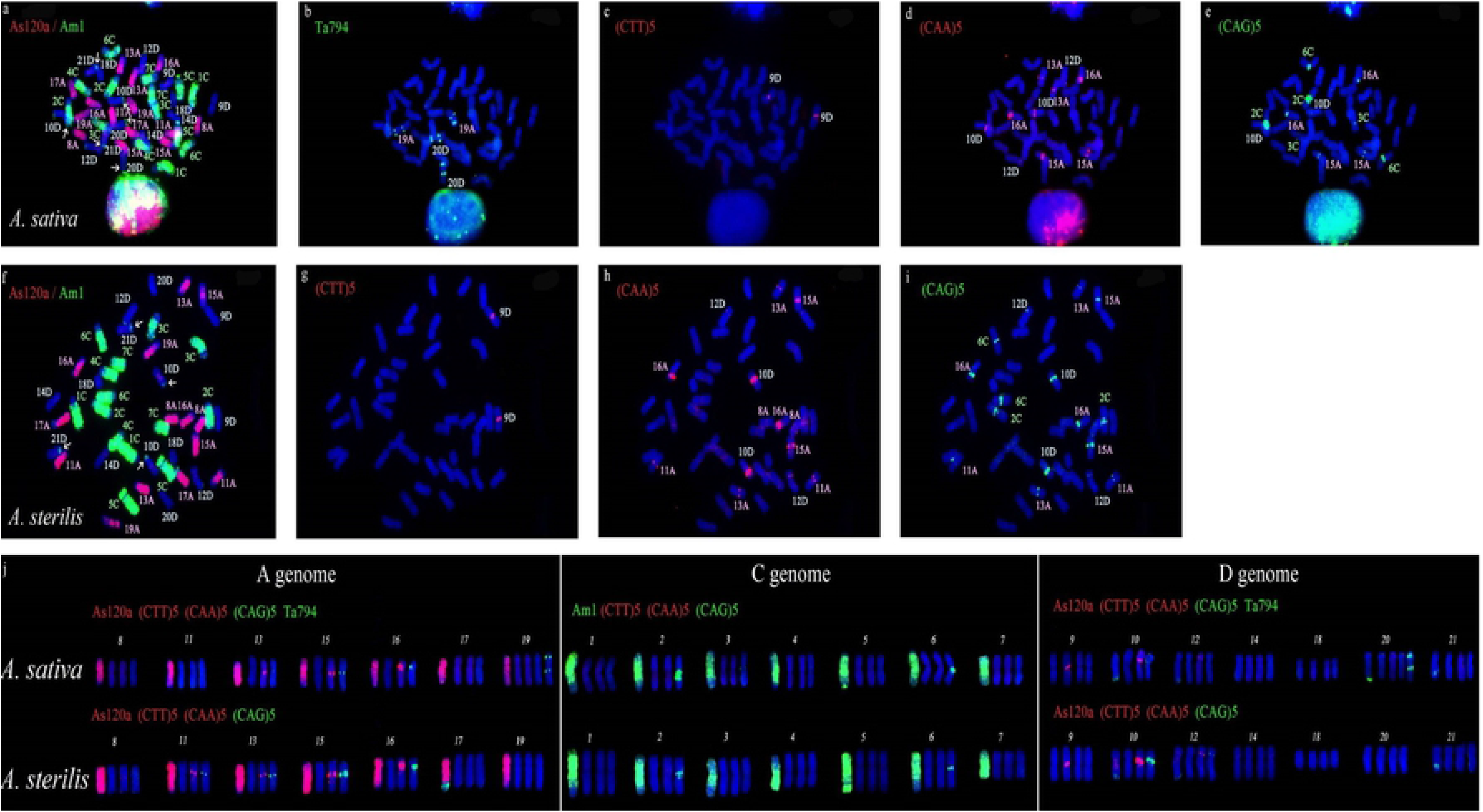
FISH performed on mitotic metaphase plates of hexaploid oats (a) *A. sativa* and (f) *A. sterilis*. (a) Simultaneous FISH of TAMRA-labeled As120a (red) and FAM-labeled Am1 (green). (b-e) The same cell as in panel ‘a’ after sequential FISH with FAM-labeled Ta794 (green), TAMRA-labeled (TTC)_5_ (red), TAMRA-labeled (AAC)_5_, and FAM-labeled (CAG)_5_ (green). (f) Simultaneous FISH of TAMRA-labeled As120a (red) and FAM-labeled Am1 (green). (g-i) The same cell as in panel ‘f’ after simultaneous FISH with TAMRA-labeled (TTC)_5_ (red), TAMRA-labeled (AAC)_5_, and FAM-labeled (CAG)_5_ (green). (j) Karyotypes showing a single chromosome of each homologous group chosen from the metaphases in panels ‘a’ and ‘f. The white arrows indicate the C/D genome translocations.

All three SSR probes produced detectable signals in the two hexaploid oats. Similar to what was found with the tetraploids, one pair of strong signals produced by the oligo-(TTC)_5_ probe was detected in the centromeric region of 9D in both hexaploids (Figs 2c, 2g and 2j). In the hexaploids, oligo-(AAC)_5_ once again produced more signals than oligo-(TTC)_5_ did. Signals from (TTC)_5_ were detected on five (13A, 15A, 16A, 10D, 12D) (Figs 2d and 2j) and six (11A, 15A, 16A, 10D, 12D) (Figs 2h and 2j) chromosomes in *A. sativa* and *A. sterilis*, respectively. All of these signals were located in centromeric or petricentromeric regions, but differed in intensity. For (CAG)_5_, the signal intensities between *A. sativa* and *A. sterilis* were similar, but differed in number. In *A. sativa*, oligo-(CAG)_5_ produced signals on five chromosomes, including 15A, 16A, 2C, 6C, and 10D (Figs 2e and 2j). All of these signals were located in centromeric regions. In A. sterilis, the number of chromosomes with hybridization signals was eight, including four A genome chromosomes (11A, 13A, 15A, 16A), two C genome chromosomes (2C and 6C), and two D genome chromosomes (10D and 12D) (Figs 2i and 2j).

## Discussion

### The chromosomal organization of SSR repeats in oat genomes

We have elucidated the chromosomal organization of three SSR motifs, (TTC)_5_, (AAC)_5_, and (CAG)_5_, in five *Avena* polyploids. All three SSR oligonucleotides produced detectable hybridization in mitotic metaphase chromosomes in these species. Most of the signals produced were located at the centrometic or petricentromeric regions (Figs 1 and 2). These results were consistent with previous studies [30–32], and implied that *Avena* genomes contain more repetitive sequences in the centromeric regions of the chromosome than in intercalary parts, as has been found in many other plants [22, 34, 38]. Signal numbers produced by these three SSR probes varied, with (AAC)_5_ producing the most hybridization signals, followed by (CAG)_5_. The (TTC)_5_ probe produced few discernable signals (Figs 1 and 2). For the tri-nucleotide repeats, A/T-rich repeats (e.g., AAG/CTT, AAC/GTT) have been shown to be predominant in dicot species [39], but that is not the case in the monocot species barley or rice. Previous studies showed that an AAT repeat gave poor hybridization signals in barley [40], and a GCC motif was dominant in the rice genome [41]. In this study, both of the A/T-rich tri-nucleotide motifs, TTC and AAC, gave poor hybridization signals in *Avena* genomes, unlike what was seen in barley and wheat, where these probes hybridized to many sites and usually appeared all along the chromosomes [29, 34]. Previous studies also showed that other tri-nucleotide motifs (AAG, TTG, ACT) hybridized poorly in *Avena* species [30–32]. Taken together, these results suggest that the tri-nucleotides, at least the A/T-rich ones, are not the predominant repeat types of SSRs in *Avena* genomes.

In *Avena* species, the C genome is highly diverged from both the A and D genomes, and the A and D genomes are of high homology [9, 42]. These differences could be observed by comparing the distribution of the SSR motifs used in this study. For instance, almost all of the signals produced by the (CAA)_5_ probe are observed in the A or D genome chromosomes, with the exception of two C genome *A. insularis* chromosomes that had detectable (CAA)_5_ signals (Figs 1s and 2j). The signal number produced by the three SSR probes also differed between the A and D genomes, with the A genome chromosomes having more signals than the D genome chromosomes (Figs 1s and 2j). The centromere plays an important role during mitosis and meiosis in higher eukaryotic organisms [38]. Centromeric sequences are the most rapidly evolved in the genome, and have been considered to generate the major differences between genomes [20]. In this study, most signals produced by the probes used were located in centromeric regions; hence, the differences in signal patterns among the A, C, and D genomes would reflect the structural differences between these genomes and support the pivotal role of the centromere in genome restructuring.

### The genomic compositions of *A. insularis, A. maroccana*, and *A. murphyi*

It is well accepted that the tetraploid species *A. insularis*, *A. maroccana*, and *A. murphyi* have been involved in the formation of hexaploid oats; however, the genomic constitutions of these species remain inconclusive. The AC genome designation was first assigned to *A. maroccana* after As and Cp genomic DNA used as probes for genomic in situ hybridization (GISH) each labeled half of the chromosomes in this species [43]. In addition, C-banding analysis showed high similarity between the chromosomes of *A. insularis*, *A. maroccana*, and *A. murphyi* [44], suggesting that they share the same genomic composition. However, these AC genome designations have been challenged by considerable evidence coming from both cytogenetic and molecular studies (summarized in Table 3). The strongest lines of evidence come from FISH analysis [17] and a GBS study [13]. An A genome-specific repetitive sequence isolated from the As diploid *A. strigosa* failed to hybridize with the genomes of the so-called AC tetraploids or the hexaploid D genome [17]. GBS markers revealed that half of the chromosomes of the tetraploids showed strong matches with the C genome chromosomes of the hexaploids, while the others showed strong matches with the D genome chromosomes [13]. In the current study, the A genome-specific probe oligo-As120a failed to hybridize with the chromosomes of the three tetraploids or the D genome chromosomes of the hexaploids (Figs 1a, 1g and 1m), once again confirming the absence of the A genome in these tetraploids. Furthermore, some C/D genome translocations were observed in *A. sativa* and *A. sterilis* (Figs 2a and 2f), and these were also detected in the three tetraploids (Figs 1b, 1h and 1n). In addition, a pair of strong (TTC)_5_ signals was detected on one A(D) chromosome in *A. insularis* and *A. maroccana* (Figs 1d, 1j and 1p). Such signals were also observed on hexaploid chromosome 9D. Together with information from previous studies, the similarities between the hexaploid D genome and the A(D) genome in all three of the tetraploids observed in this study provide robust evidence for the presence of the D genome in these tetraploids, strongly suggesting that the three tetraploid species in the section *Pachycarpa* should be re-designated as having CD genomes, rather than AC genomes.

**Table 3.** Evidence for the DC genome assignment of the tetraploid species in the section *Pachycarpa*

| Species | FISH | Molecular marker | Chromosome pairing behaviour |
| --- | --- | --- | --- |
| *A. insularis* | Linares et al. [17] | Yan et al. [13] | Loskutov [45] |
| *A. maroccana* | Fominaya et al. [33];<br>Linares et al. [17] | Oliver et al. [46, 47];<br>Chew et al. [16];<br>Yan et al. [13, 18] | Ladizinsky [48] |
| *A. murphyi* | Linares et al. [17] | Peng et al. [49-51];<br>Yan et al. [13, 18] |  |

### The tetraploid progenitor of hexaploid oats

No conclusive agreement has been reached regarding which tetraploid species may have contributed to the hexaploid oat genome. Examining the existing literature, all of the three tetraploid species in the section *Pachycarpa* have been postulated to be the tetraploid ancestor of the hexaploids at one time or another [10, 13–15, 18]. In the current study, the signal patterns produced by the probes used revealed that the three tetraploids were closely related, but well differentiated from each other. For instance, the number of intergenomic translocations in *A. murphyi* (Fig 1n) differed from that in *A. insularis* (Fig 1b) and *A. maroccana* (Fig 1h), while the signal patterns produced by (TCC)_5_ allowed for the differentiation of *A. maroccana* (Fig 1j) from the other two tetraploids (Figs 1d and 1p). However, none of these tetraploids showed a FISH karyotype that is better matched to the hexaploids than the others in this study. One possibility is that it wasn’t one of the extant tetraploids that participated in the formation of the hexaploid oats, but, rather, a common ancestor that has not been identified or is now extinct. There is considerable evidence that all three tetraploids originated from the same tetraploid ancestor and then diverged from one another after several large chromosomal rearrangements and other changes in their chromosomes decreased their level of homology [44, 52, 53]. Another plausible explanation is that the genomes of the hexaploids may have experienced substantial restructuring after polyploidy took place. This hypothesis is supported by previous study, which showed significant genome downsizing after poyploidizations in genus *Avena* [54]. A similar phenomenon has been observed in wheat. Zhang et al. [55] observed distinct differences in multiple phenotypic traits and identified a large number of differentially expressed genes between the natural AB genome tetraploid wheat and a ploidy-reversed (from hexaploid to tetraploid) “extracted tetraploid wheat” which has AB genomes that are virtually identical to the AB sub-genomes of its bread wheat donor. In hexaploid oat, there exists a genetic mechanism that is similar to the *Ph1* locus in hexaploid wheat, which ensures exclusive homologous chromosome paring in meiosis [56]. This attribute of hexaploid oat would make possible the reconstitution of its CD component by a simple backcrossing technique, and, therefore, could provide a unique opportunity to address whether and to what extend the CD component of hexaploid oat has been modified during its evolutionary history at the allohexaploid level, and provide more substantial evidence on the tetraploid ancestor of hexaploids.

### Conclusions

FISH techniques with six repetitive DNA sequences showed a distinct hybridization patterns of *Avena* A, C and D genomes, confirming the substantial structural differences among these genomes, particular the large divergence between the C and A/D genomes. Several intergenomic translocations between the D and C genomes in hexaploid oats were detected, and such intergenomic translocations were also observed in all three tetraploids in the section *Pachycarpa*, providing good evidence for the presence of the D genome in these tetraploids, hence supporting a final re-designation of these tetraploids as CD genomes.

## Acknowledgments

We gratefully acknowledge the fellows in Triticeae Research Institute who gave us professional and technical assistance during the experiment, and we thank Dr. Eric N. Jellen, Brigham Young University, Provo, Utah, USA, for kindly providing the seeds of *A. insularis*.

